# The historical range and drivers of decline of the Tapanuli orangutan

**DOI:** 10.1101/2020.08.11.246058

**Authors:** Erik Meijaard, Safwanah Ni’matullah, Rona Dennis, Julie Sherman, Onrizal, Serge A. Wich

**Author notes:** Corresponding author, (EM).

## Abstract

The Tapanuli Orangutan (*Pongo tapanuliensis*) is the most threatened great ape species in the world. It is restricted to an area of about 1,000 km^2^ of upland forest where fewer than 800 animals survive in three declining subpopulations. Through a historical ecology approach involving analysis of newspaper, journals, books and museum records from the early 1800s to 2009, we demonstrate that historically *Pongo tapanuliensis* inhabited a much larger area, and across a much wider range of habitat types than now. Its current Extent of Occurrence is between 2.5% and 5.0% of the historical range in the 1890s and 1940s respectively. A combination of historical fragmentation of forest habitats, mostly for small-scale agriculture, and unsustainable hunting likely drove various populations to the south, east and west of the current population to extinction. This happened prior to the industrial-scale forest conversion that started in the 1970s. Our findings indicate how sensitive *P. tapanuliensis* is to the combined effects of habitat fragmentation and unsustainable take-off rates. Saving this species will require prevention of any further fragmentation and killings or other removal of animals from the remaining population. Without concerted action to achieve this, the remaining populations of *P. tapanuliensis* are doomed to become extinct within several orangutan generations.

## Introduction

Determining the key drivers of population decline is a primary objective in conservation biology and wildlife management. Many wildlife species are threatened by a range of different and often interacting factors, and developing effective conservation strategies requires unravelling how these threats interact [1]. This is rarely easy, because species operate in complex socio-ecological systems in which different components are affected by a range of anthropomorphic factors such as habitat loss and fragmentation or unsustainable harvest. Evidence-based conservation seeks to address this by quantifying the relationships between conservation actions, change in threat severity and change in conservation status [2, 3]. Collecting evidence is, however, time-consuming, and when conservation problems are “wicked”, i.e., the problems change as solutions are found [4], a stable solution may not be found to a particular conservation problem [5]. This often means that scientific evidence does not support clear-cut conclusions in value-driven debates that characterize conservation [6]. Nevertheless, conservation advocates often seek simple narratives to convince the public of the urgency of environmental problems and the need to support it.

One way to bring more clarity in often polarized debates around simple narratives is to be more specific about the system in which a particular problem plays out. For example, if the system boundaries are limited to oil palm as an ecological threat to orangutan survival [7], a simple solution would be to ban palm oil use and to stop its production, preventing further deforestation. If the system boundaries are extended to include smallholder farmers who produce palm oil for their own needs as well as international markets, a ban on palm oil would encompass broader ethical connotations as it would affect people’s livelihoods [8]. The use of different perspectives in complex conservation contexts may not make it easier to solve them but can provide helpful insights about the system boundaries of a particular problem. Are they, for example, mostly ecological, or do they involve human threats, such as hunting, or societal ethics? One such perspective is history. Looking back in time on the development of a particular problem may provide insights about the underlying drivers of that problem [9]. The historical ecology approach uses historical knowledge on the management of ecosystems or species [10]. Referring to historical evidence has, for example, provided valuable understanding about the ecology of orangutans and what likely caused their decline during the Late Pleistocene and Holocene, which informs their management today [11]. Here we apply an analysis of historical ecology to one particular species of orangutan, *P. tapanuliensis*, by analysing rarely used colonial-era literature to better understand the historical distribution range of the species and the different ecological conditions (elevation, vegetation types) in which it occurred. Indonesia’s colonial literature on natural history was mostly written in Dutch and German, and is not commonly used by conservation scientists working in Indonesia.

*Pongo tapanuliensis* was described in 2017 as a third species of orangutan [12], 20 years after this orangutan population was formally reported to modern science [13]. The species is restricted to three areas of mostly upland forest in the Batang Toru area in North Sumatra (Fig 1), totalling approximately 1,023 km^2^ [14, 15]. This orangutan population had been largely overlooked by science, despite having been tentatively described in the colonial literature [16]. The estimated total number of wild *P. tapanuliensis* is currently 767 [95%: 213-1,597, 14] making this the great ape species with the lowest number of individuals in the wild and perhaps the most threatened in the world [17].

**Fig 1.**
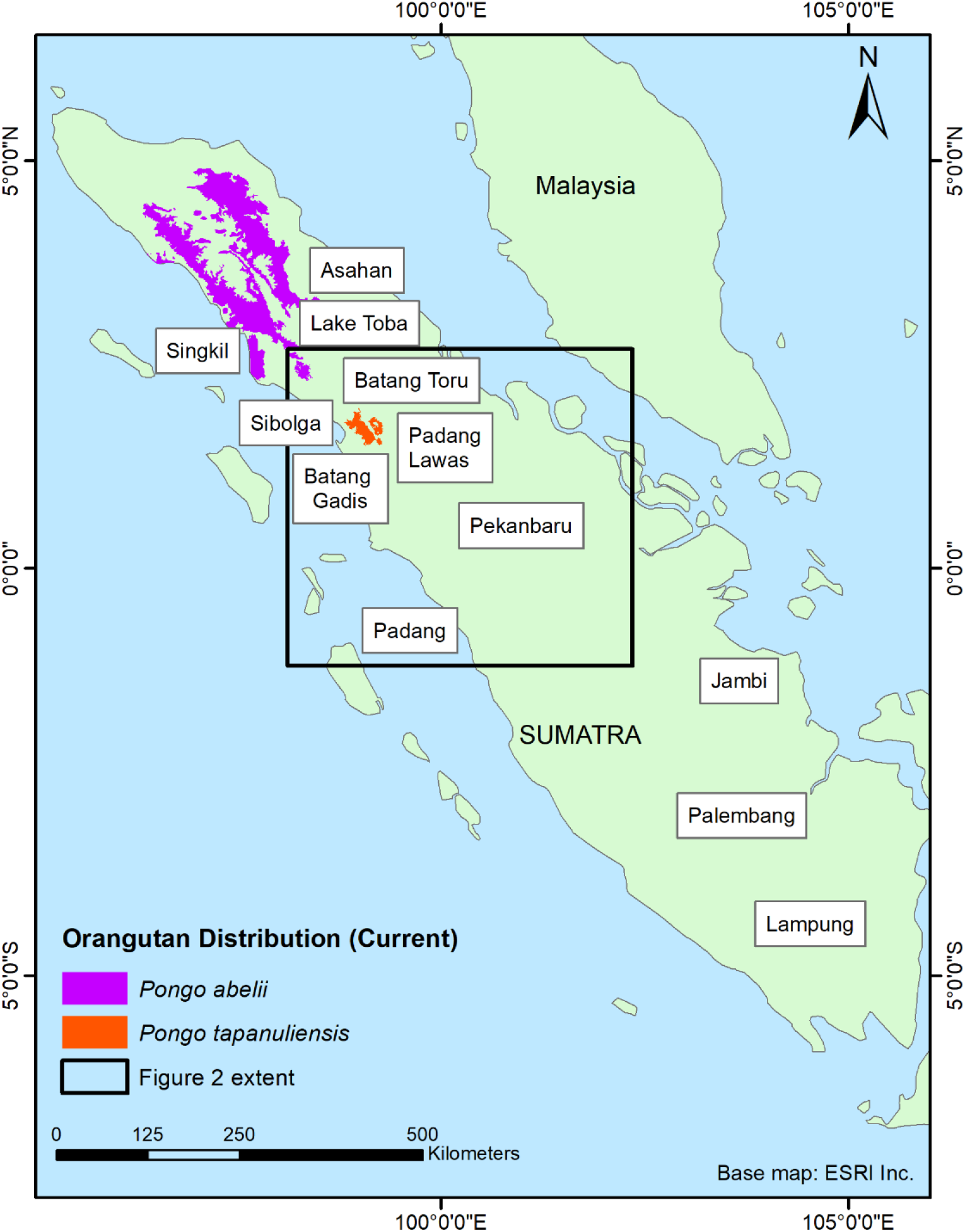
Map of the island of Sumatra, showing the current distribution ranges of *P. tapanuliensis* and *P. abelii*, and the main locations (cities, districts and other geographic features) mentioned in the text. The inset shows the area of Fig 2. The inset shows the area of Fig 2. Reprinted from [18] under a CC BY license, with permission from ESRI, original copyright 2000.

The species is currently under threat of habitat loss from agriculture, hunting and conflict killing, and development in the area for infrastructure, gold mining, and geothermal and hydro-energy. These threaten to further reduce and fragment remaining habitat, reduce dispersal opportunities for the orangutans between subpopulations, and undermine population viability through unsustainable mortality rates [14, 19-22]. Due to its restricted current distribution mostly centred around higher elevations (834.4±219.3m asl) compared to 701.7±454.8 m asl for the Sumatran orangutan (*P. abelii*) and 170.6±187.0 m asl for the Bornean orangutan (*P. pygmaeus*) [23], it has been argued that the individuals of the species have adapted specifically to the uplands that cover most of its current distribution in Batang Toru [19]. What is not clear is whether the current existing altitudinal differences between the orangutan species are the result of ecological specializations to highland ecological conditions, or whether the highland species now occur at higher elevations because their previous lowland habitats no longer exist or because the species became extinct there. The fossil record for Sumatran orangutans confirms that the genus *Pongo* was once more widespread. Extensive remains from the Late Pleistocene and Holocene have been excavated from a range of caves in the Padang Highlands, some 300 km south of the current range [24] (Fig 1). Why the species disappeared from that part of Sumatra remains unclear but unsustainable hunting is one of the possible explanations, because until recently large areas of suitable forest habitat remained in areas where orangutans are now extinct [25]. Given that, in the past, forest cover was also much more widespread in the range of *P. tapanuliensis*, it is important to determine whether historically (ca. past 500 years) orangutans did occur in those areas. This would help establish whether *P. tapanuliensis* has indeed evolved to only live in the highlands and to estimate what its past distribution could have been.

The aim of this paper is to compile reports of orangutans occurring to the south of Lake Toba (Fig 1) with the focus on determining how reliable these are, and, where feasible, provide a location for the occurrence of orangutans to assess whether these are predominantly highland sites, and to assess which factors could have led to their disappearance in those areas. Based on the information we develop historical distribution maps as reference points for estimating historical population declines and their drivers, and better understand the ecological conditions under which the species used to occur. With this information we seek to inform current conservation strategies and provide data to conservation practitioners for setting long-term recovery targets for the species to ensure full ecological functionality [9].

## Methods

We compiled records of orangutans from historical sources by searching natural history books, scientific papers, and historical newspapers from before 1940. We searched databases, including the Biodiversity Heritage Library (https://www.biodiversitylibrary.org/) and online historic newspapers, books and journals (https://www.delpher.nl) with location specific keywords such as Sumatra, Batang Toeroe, and Tapanoeli, using Dutch spelling. We combined this with searches for terms specifically referring to orangutans: Orang oetan, orang-oetan, orangutan, and also mawas, mias and maias (local names for orangutan commonly used in historical literature), using a variety of spellings. For the period since 1940, we used the sources from the review in Rijksen and Meijaard [25] as well as scientific papers and personal communications. To determine the locations of the historical sightings or captures we consulted the online Leiden University Library colonial map repository (http://maps.library.leiden.edu/apps/s7). In some cases, rivers or villages were indicated which made it feasible to estimate the location of the sightings quite accurately. In other cases, the area of the sighting or captures was indicated in a broader area which reduced accuracy (tens of kilometres).

We assessed the likely vegetation types that *P. tapanuliensis* would have occurred in, and determined the elevation at which they were reported. For this, we vectorised a high-resolution scanned copy of the first official forest cover map of Indonesia [26], dated 1950 and at a scale of 1:250,000,000, which is likely based on maps produced by the Netherlands-Indies cartographic service from the 1930s and 1940s. In order to analyse the map and integrate with other spatial layers in a GIS, we automatically vectorised the map using the ArcScan extension within ESRI ArcGIS [27]. The first step in the process was to geo-reference the scanned map to the coastal boundary of Sumatra. The next step was extraction of the area of interest which was then vectorized. This resulted in numerous polylines which were cleaned and edited to produce polygons representing various land cover classes. The last stage in preparing the 1950s map for integration with other spatial datasets in the GIS was to eliminate spatial distortion as much as possible. Old hand-drawn maps, which in this case was an Indonesia-wide map, have inherent distortion when compared to modern maps. We used a process called rubber-sheeting to make small spatial adjustments in the vectorized georeferenced map to align individual parts of the map more accurately with the coastline and inland features such as lakes and rivers. Although not a perfect match, we are satisfied with the spatial accuracy of the vectorized 1950s map in terms of meeting the objectives of this analysis.

While the exact location of the historical orangutan sightings cannot be determined with certainty, the descriptions often provide sufficient detail through names of rivers and villages to estimate the elevation and dominant vegetation where they occurred. Elevation was determined from the altitude layer in Google Earth Pro [28]. We used the vegetation map for Sumatra [29] in combination with knowledge gained by co-author SW during surveys in the region to assign one of the forest categories to an estimated historical location.

To approximate the population decline of *P. tapanuliensis* in historical times, we mapped the historical range. This provides important insights about the areas and vegetation types that the species once occurred in, providing insights regarding its ecological functionality [30]. Grace et al. [9] recommend using either a 1500 AD or 1750 AD target year for mapping the historical range, unless the historical data are insufficiently accurate or reliable. Because there is uncertainty about all historical species data (unless supported by specimens), we developed two maps: 1890s and 1940s. Each of these periods have different data sources with varying reliability associated with them. By presenting these different maps, conservation scientists and policy-makers can debate the merit of accepting either of these two (or a different map altogether) to set a historical baseline for the species. Historical ranges were mapped by modelling watersheds containing historic orangutan records of breeding females separated by large rivers. We divided the island of Sumatra into potential orangutan subpopulation ranges by mapping large rivers [based on 31]. These boundaries were chosen with the assumption that female orangutans rarely disperse across these boundaries into a neighbouring watershed. Whereas subadult and adult males do disperse through areas of higher elevation and low-quality habitat, females are very rarely seen in such locations [25, 32-34]. This lack of inter-watershed dispersal is supported by genetic studies [33, 35]. We assume that if one of our subpopulations is reduced below carrying capacity or goes extinct, the probability of recolonization by immigrating females is negligible. An overlay of the historical distribution range (pre-commercial timber industry) and the potential subpopulations ranges resulted in a map of areas with or without orangutan populations. To further enhance the analysis, we also evaluated which historical points were located above or below the 750m elevation contour to assess whether or not there was a greater number of observations at higher elevations in the historic records. We accessed the NASA Shuttle Radar Topography Mission Global 1 arc second data from the NASA EOSDIS Land Processes DAAC to download the digital elevation model tiles for Sumatra from which we extract the 750 m contour information [36].

## Results

### Historical accounts

We report the various historical accounts of orangutan sightings or specimens from outside the currently known range in chronological order, starting with Nikolaas Tulp [37] who in 1641 reported on a specimen of “Indian Satyr” he had received, which had been collected in “Angola”, most likely the former district of Angkola (no. 1 in Fig 2) [25], which is now part of South Tapanuli District. His descriptions and drawings indicate an orangutan, “of female sex”, as Tulp writes. Several authors [e.g., 38, 39] argued that Tulp more likely referred to the African country of Angola and that his specimen was therefore likely a chimpanzee or gorilla and not an orangutan. Rijksen and Meijaard [25], however, pointed out that Tulp specifically referred to his “Indian” specimen being distinct from the African species, and also mentions that the species occurs of the island of Borneo.

**Fig 2.**
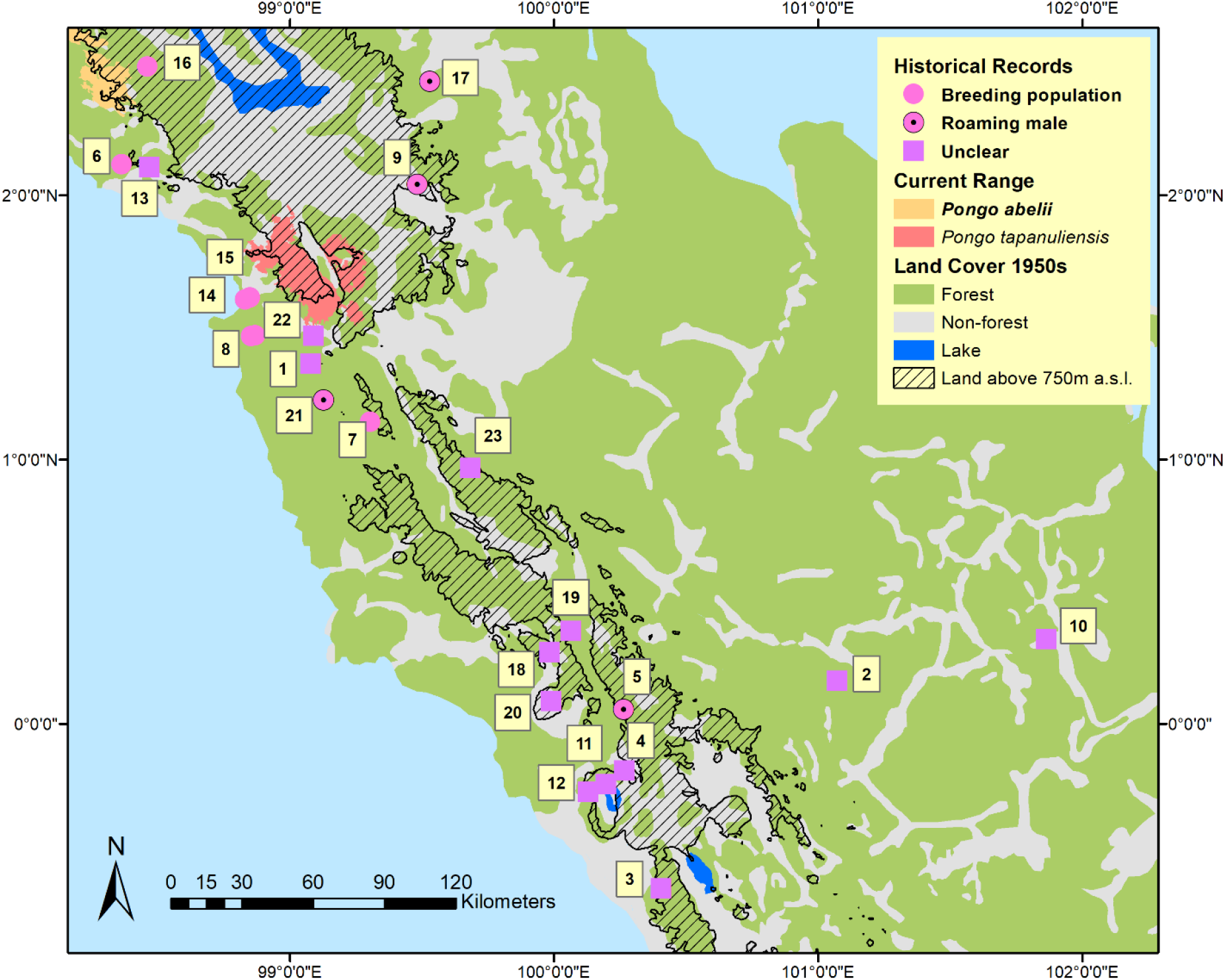
Map showing points where orangutans were historically reported, land above 750m, and the forest cover in 1950 before large-scale commercial forest exploitation began. Reprinted from [26] under a CC BY license, with permission from PT Balai Pustaka, original copyright 1950.

Schlegel and Mueller [40] reported in 1841 on two orangutan crania that were obtained by a military doctor in the environs of Jambi, some 650 km south-east of the current closest orangutan population (Fig 1). It is unclear whether these animals were obtained in Jambi from the wild or whether they were in captivity and possibly originated from northern Sumatra. Schlegel and Mueller reported that the two skulls were housed in the “Rijks-Museum”, which presumably refers to the Leiden Museum of Natural History. We were unable to locate these specimens in the Leiden collection. Schlegel and Mueller further wrote that on the west coast of Sumatra, especially north of the equator, the orangutan was known by the name *mawej*, although in areas further south such as Indrapura and Bengkulu, the names *orang-panda* or *orang-pandak* were used. Co-author Onrizal, remembers growing up east of Padang (Sungai Dareh) listening to stories about *orang-pendek*, human-like creatures living in the forest, which reportedly ceased to exist in the area in the 1970s. Stories of such *orang-pendek* [or other names, such as gugu, sedapa, orang lètje or orang segagang, see 41,42] abound in other parts of southern Sumatra and there has been speculation for over a century that these could be remnant orangutan populations [43-46], although evidence remains lacking. For the purpose of the current paper we do not focus on the *orang-pendek* narratives, but recognize that many of these narratives could indeed refer to orangutans, as suggested compellingly by Forth [42].

Schlegel and Mueller [40] acknowledged that the orangutan was especially common in the north-east of Sumatra, but that occasionally they were encountered further south and along the western shores of Sumatra. Also, the species had been reported from Indrapura (in present-day Riau Province) and near Padang (in West Sumatra, Fig 1), although the descriptions by Schlegel and Müller are insufficient to confirm that these reports referred to orangutans. Earlier writings, in 1837, by Müller [47], however, clarify this, as he refers to “orang-oetan” which locals named “mawej” that one could occasionally encounter in the extensive forests and swamp forests in, what is now, Kampar Regency, roughly between the towns of Salo and Gunung Sahilan (no. 2 in Fig 2).

A letter from an anonymous Dutch missionary published in 1892 [48] mentions orangutan sightings, although the letter was 50 years old then, so approximately referring to 1842. It mentions that “monkeys, and especially ‘orang-oetangs’” made life difficult to people travelling inland from Padang, by throwing “stones, coconuts, branches, and others” at travellers through the “Padansche Bovenlanden”, approximately in the area that is now Bukittinggi (no. 3 in Fig 2). It specifically mentions an event during which someone was attacked by orangutans half-way between Fort de Kock (in Bukittinggi) and Bonjol (no. 4 in Fig 2).

In further description of their travels across Sumatra, Müller and Horner [49] wrote in 1855 that orangutans were not unknown in the Tapanuli area and especially common in “Taroemon”, i.e., present-day Trumon in the Singkil area (Fig 1), which is part of the *P. abelii* range. They report that people distinguished between two types of orangutans, the *maweh baroet* (baroet meaning monkey in the local language) and *maweh orang* (the ‘human’ orangutan). Ludeking [50], in his descriptions of West Sumatra, mentions, in 1862, a record of a six feet tall, upright primate, possibly an adult male orangutan, that his informants had seen on *Bukit Gedang*, which is close to present-day Bonjol (no. 5 in Fig 2). A decade later, in 1878, von Rosenberg [51] did not provide much detail but similarly mentioned that orangutans were present north of “Tapanoeli” (what is now Sibolga) to Singkil (Fig 1), indicating presence of the species in the coastal lowlands west of Lake Toba (no. 6 in Fig 2). He saw two orangutans but did not clarify where he saw them, although Snelleman [52] stated that von Rosenberg had seen “two youthful specimens” in the area between Tapanuli and Singkil.

In 1879, Kramm [53] reported on a hunting expedition near “Soeroe Mantinggi”, where he found several orangutans and observed them for several hours. The location likely referred to Sayur Matinggi (no. 7 in Fig 2), which is currently located in the Batang Gadis area, some 50 km south of the current range of *P. tapanuliensis*. Kramm mentioned that Soeroe Mantinggi is located at a distance of 22 *“palen”* from Padang Sidempoean. A “paal” was a measurement used in the Netherlands-Indies, equalling 1852 m on Sumatra, indicating a distance of about 40 km for 22 *“palen”*. Sayur Matinggi is currently located some 26 km from Padangsidempuan, which indicates that indeed this is likely to be the location where Kramm observed orangutans. Kramm was familiar with orangutans which he reported to have also encountered in “Loemoet” and “Batang-Taro”. We believe that the former refers to Lumut (no. 8 in Fig 2), just south of Sibolga (Fig 1), and that Batang-Taro is an older name for the Batang Toru area, where *P. tapanuliensis* occurs until today.

Orangutans also seem to have occurred northeast of the current range of the *P tapanuliensis*. In 1885, Neumann [54] described the species from “Hadjoran”, which was located in the watershed of the “Batang Si Ombal” and “Aek Hiloeng”, and for which the following coordinates were given: N 2°1’25” and E 99°29’, in the current district of Padang Lawas (Fig 1, no. 9 in Fig 2). This is about 50 km northeast of the most eastern current range of *P. tapanuliensis*. The detailed description, however, suggests that the species was very rare there, and the people of “Hadjoran” had not seen the orangutan there before. The animal was shot, with descriptions of the local people suggesting it was at least 1 m tall, possibly indicating an adult male, which are known to roam far from breeding populations. Neumann writes that he travelled extensively through forest areas in the Padang Lawas area searching for orangutans but never managed to encounter one.

In 1887, Snelleman [52] mentioned a report from a government employee in Lampung (Fig 1) who had heard about orangutans in that part of far southern Sumatra. Several hunts were organized to find the orangutans but these were unsuccessful, and also when local people were asked they stated that they that had only heard about orangutans in that area from hearsay but that they could not pinpoint where orangutans were supposed to occur. We did not add these records to the maps as their reliability seems low.

An interesting reference to orangutans, as far south-east as Pelalawan is provided by Twiss [55] who, in 1890, described a record reported in the Tijdschrift van het Binnenlandsch Bestuur, Vol. III, p. 138 by mr “L.H.” [56], who was on a trip to *Poeloe Lawan* (Pelelawan) near the confluence of the Batang Nilo and Kampar Rivers (no. 10 in Fig 2) in 1888, and had an orangutan in his visor, but decided not to shoot it as he had nothing on him to prepare the skin. Twiss [55] reported that L.H. was familiar with orangutans from ones he had seen on Borneo, so the chance of a mistaken identification are small. Twiss also reported that 18 to 20 years prior to his writings (i.e., around 1870), an orangutan was shot in the mountains around Lake Maninjau (no. 11 in Fig 2), while older people remember seeing orangutans, albeit very rarely, in forests on Bukit Silajang (no. 12 in Fig 2), a mountain near Lubuk Basung [55].

In 1890, Hagen [57, p. 66] stated that orangutans were known from the west coast between Tapanuli and Singkil (no. 13 in Fig 2), although Singkil is in the range of *P. abelii* and it is not clear whether the coastal Tapanuli reference referred to the area of the current range of *P. tapanuliensis*, or whether it referred to what is now the Central Tapanuli District which extends to the southern part of Singkil, west of Lake Toba. Interestingly, he referred to an orangutan from the interior of Padang (in reference to an article by S. Jentink, Aardrijkskundig Weekblad, 1881, No. 44, p. 287 – not seen by the authors), in west Sumatra that ended up in the

Rotterdam Zoo where it died of a bone deformation disease (the skull is kept in the Natural History museum in Leiden, the Netherlands: RMNH.MAM.544).

Miller [58, p. 483] in his account about the mammals collected by W.L. Abbott on the west coast of Sumatra in 1901 and 1902 mentioned the following about orangutans: “The orang utan exists, but not abundantly, about Tapanuli Bay. Two miles up the Jaga Jaga River (no. 14 in Fig 2) some nibong palms were seen that had been broken off by orangs, and also an old sarong (shelter), but the traces were old. There were said to be more a few miles farther inland, particularly up the Berdiri River (no. 15 in Fig 2). The natives say they always go about in pairs.” Miller described the Jaga Jaga River as “a stream near the south end of the Tapanuli or Sibolga Bay”. We located the Berdiri River on an old map under the name “Bardari River”, and we located “Djaga Djaga” as well.

In 1904, Beccari [59] reported orangutans around Rambung, in the Tapanuli region, and in the hinterland of Sibolga, where he collected a specimen. We were unable to determine the location of Rambung but there is a Rambong north of the Singkil area, and thus well in *P. abelli* range. The hinterland of Sibolga could either refer to the current *P. tapanuliensis* population or the historical range - we were unable to determine whether this specimen still exists, and, if so, where. Beccari further stated that “in the Zoological Museum at Florence is the skeleton of a young orang-utan, described as coming from Palembang (Fig 1), on the east coast of Sumatra”, some 800 km southeast of the nearest current orangutan population. We contacted the curator of the Florence museum, who wrote in response that the specimen was indeed present under specimen number MZUF-12: “The specimen was purchased in 1889 in London (G.A. Frank, 9 Haverstock Hill, London). It is a subadult male. The place of origin is Palembang, but it may have been captured elsewhere. There are no manifest connections with O. Beccari.” Gustav Adolf Frank was a well-known natural history trader based in Amsterdam and London, and he probably had a good network of local suppliers. The description of the skeleton provided by Agnelli and colleagues [60] is inconclusive as to what species it belongs to. We can therefore not know for sure whether the animal was caught near Palembang and transported to Europe from there, or whether it originated from northern Sumatra (either of the two known species).

Also in 1904, Volz [61] wrote about the distribution of orangutans on Sumatra, although it is not clear to what extent the information is informed by Volz’ own surveys or interpretation of secondary information. Volz suggested that there were no orangutans east of the Langkat River, which he thought was likely the remnant of a large bay or sea connection that once separated north and south Sumatra approximately in a line from Sibolga to north of Medan. He expanded on this in his work a few years later [62], in which he also described additional orangutan sightings. This included a sighting in the upland area west of Lake Toba at an elevation of 1,400 m asl (no. 16 in Fig 2). Referring to the same area, Kohler [63] described, in 1926, a visit to Sibolga where the host had a young orangutan which had been caught in the forest on the west of Lake Toba, indicating a breeding population there. Volz [61] also described a sighting of an orangutan east of Lake Toba in the upper Kualu River area (ca. N 2°26’ E 99°32’; no. 17 in Fig 2). Again, however, the description of a large ape that moved ‘slowly and ponderously’ may suggest an adult male, and because people there are not familiar with the species, possibly a wandering male outside the range of a breeding population.

In a 1930 article, Delmont [64] described a hunting expedition on the upper Musi River, near “Sekajoe” in the foothills of the Barisan mountain range in what is now South Sumatra Province. His informant, Mr Ghoba Ramah, told him that orangutans were particularly common in the area and were raiding the crops of local farmers. After four weeks, they managed to catch seven orangutans. They then moved to a location four hours rowing upstream, where they quickly observed a female orangutan with young. They set out cages with fruit bait for capturing orangutans, but the first morning after arrival they had only managed to catch some monkeys and a pig. After that they were more successful and claimed to have caught one male orangutan and a female with young, and over the next few days they caught several more orangutans. Delmont’s stories are intriguing but strike us as somewhat fantastical, as it is unlikely that anyone could catch significant numbers of orangutans with baited cages. More likely these could be Pig-tailed Macaques, *Macaca nemestrina*, who indeed move about in groups, raid crops and can be trapped in cages. We therefore do not consider this source to be reliable, and do not include this record in Fig 2 unless evidence (e.g., specimens) is found for Delmont’s claims in 1935.

The various historical accounts above were summarized in 1935 in a map drawn by van Heurn [65] which shows that clearly the conservation community was aware of the existence of orangutans south, west and east of Lake Toba. Interestingly, though this map depicts the current Batang Toru population to be part of the range where the species had become extinct, while the only extant population is a narrow band to the east of Lake Toba in the Asahan District (Fig 1), where the species is not currently known. It suggests that information about orangutan distribution was still rudimentary in the 1930s, which may have the reason for a request to C.R. Carpenter [66] who conducted a survey on behalf of the Nederlandsch-Indische Vereeniging tot Natuurbescherming. He worked mostly in the northern parts of Sumatra and sent questionnaires to Dutch soldiers stationed in areas where orangutans could potentially occur. Carpenter assumed that orangutan did not occur south of a line drawn from Singkil to the Sumatran east coast, thus overlooking much of the historical evidence of orangutans south of Lake Toba. Carpenter’s questionnaires, however, included three reports of orangutans outside the known range. The first is from Captain H.J. Kloprogge who had been based in Aceh, Siak, Indrapura and Pekanbaru (Fig 1) between 1921 and 1936, spending an average 12 days per months in the forest on patrol. He claimed to have seen orangutans “two to four” times during forest patrols, and indicated their presence on the hand-drawn map accompanying the questionnaire throughout Aceh and the current Batang Toru range area. Second, Captain M. Kooistra reported seeing 12 orangutans in Aceh and also indicated them as present on his map near Jambi (Muara Tembesi), where he had been stationed in 1925 and 1926 (Fig 1). As there is no further information about this record, we do not include it in Fig 2. Third, Captain H.G.C. Pel was stationed in Siak (near Pekanbaru, Fig 1) from 1933 to 1935, and reported seeing an orangutan in captivity north of the town of Talu (no. 18 in Fig 2), on a tributary of the Kampar River. We consider it unlikely that such a captive orangutan in a remote village would have been transported to the area, and map this point as likely present. We do not map a report of orangutan presence in Batang Toru and Sipirok from 1939 [67], as these are still part of the current range.

There seems to be a gap in records between the 1930s and 1970s, but in the early 1970s, the Indonesian forester Kiras S. Depari reported orangutan sightings along the Batang Toru River, in the Sibual-buali Reserve and in the Rimbu Panti Wildlife Reserve (no. 19 in Fig 2) [16]. In 1976, Borner [68] also noted that a 10-12 year old male orangutan had been shot just outside Rimbu Panti, and that villagers had shortly before seen “two other black ‘orang-utans’ walking on the ground”, but his surveys could not find any nests. Borner also interviewed timber workers in Torgamba in the South Labuhan Batu District, who said that orangutans occurred in those forests but Borner’s surveys could again not verify this. Two other primatologists, C.C. Wilson and W.L. Wilson [69, not seen] confirmed the presence of orangutans in South Tapanuli in 1977, while also reporting them around Pekanbaru, in Riau Province. Herman Rijksen (*in litt*., 17 Oct 2019) reported receiving two captive juvenile orangutans, which, according to the Indonesian conservation authorities, had been confiscated in “Angkola”. Finally, the presence of orangutans was indicated by a botanist and a wildlife researcher on Gunung Talamau in the late 1980s (no. 20 in Fig 2; Laumonier, pers. comm.). Presence in this region was confirmed by a survey in 1996 by Rijksen, Meijaard and van Schaik [13], when several nests were found on the edge of this Reserve, but follow up surveys by SW could not confirm this report and suggested that the nests may have been eagle nests. Based on surveys in 1997, Meijaard [13] did describe various reported orangutan sightings from the Batang Gadis area, including a large, possibly male, orangutan close to the Bhara Induk logging base camp (no. 21 in Fig 2), although field surveys did not reveal any nests.

More recently, Wich et al. [70] found several orangutan nests in the peat swamp forests near Lumut (no. 8 in Fig 2) and heard a male long call in the same area. Local community members mentioned that they had also seen orangutans in the area [70]. Approximately 2 km south of the Batang Toru River (southeast of the village of Batang Toru), a geologist (Martin Jones) spotted a solitary orangutan in the forest in 2004 (no. 22 in Fig 2). Finally, Bradshaw, Isagi [71] reported on orangutans in the Barumun Wildlife Reserve in the Padang Lawas District (no. 23 in Fig 2). Nests were reportedly encountered and one staff of the local conservation department reported a direct encounter with an orangutan in 2009.

While there is significant spatial inaccuracy in the historical records of *P. tapanuliensis* outside the current range, we can still make an educated guess of the different habitats and elevations in which these populations occurred (Table 1). Habitats in which the species once occurred included tall peat swamp forest, freshwater swamp forest mosaic and secondary forest, forest on limestone, hill forest between 300 and 1,000 a.s.l., and submontane forest between 1,000 and 1,800 m a.s.l., indicating the full range of habitats that is also used by *P. abelii* [16].

**Table 1.**
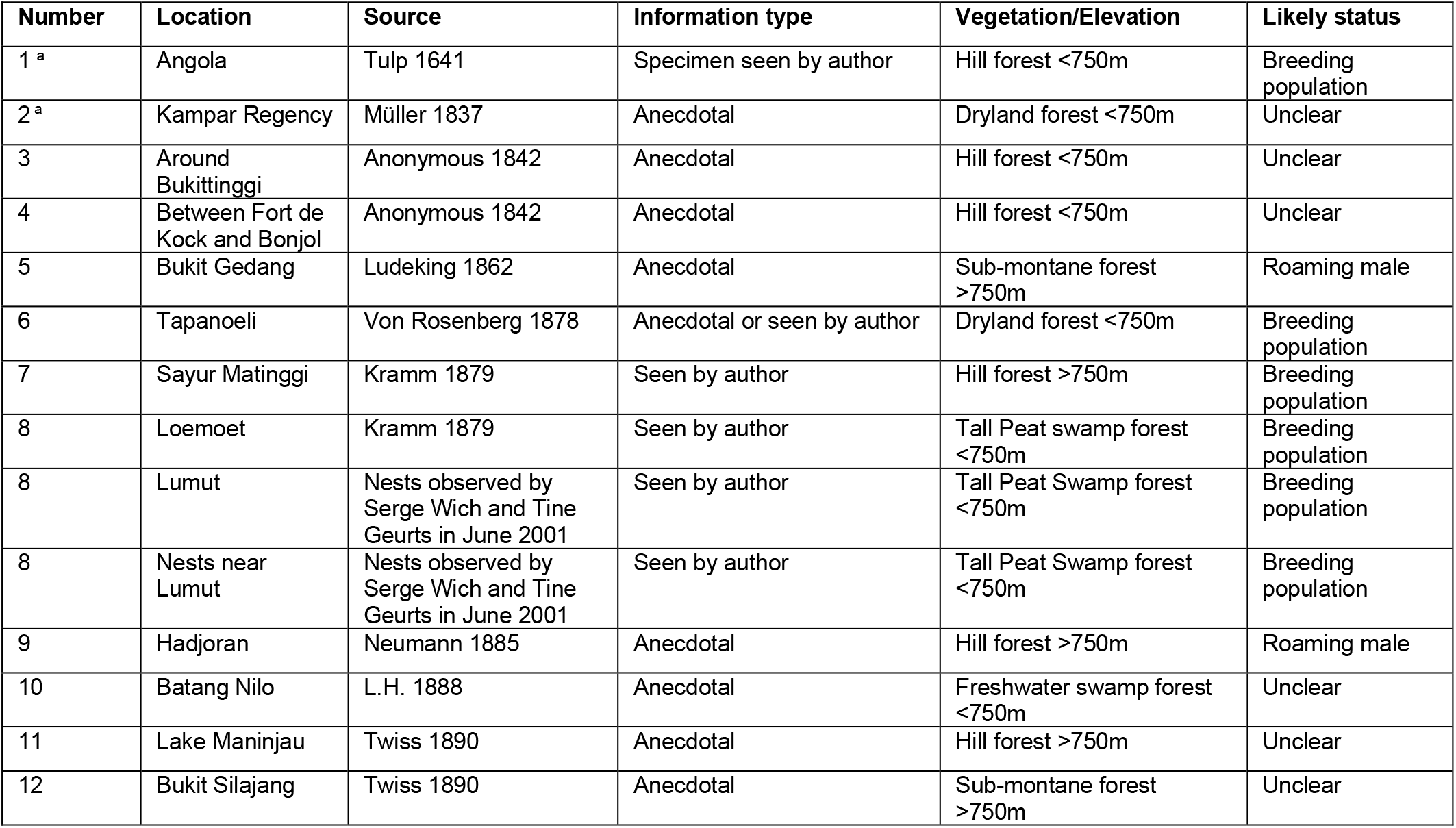
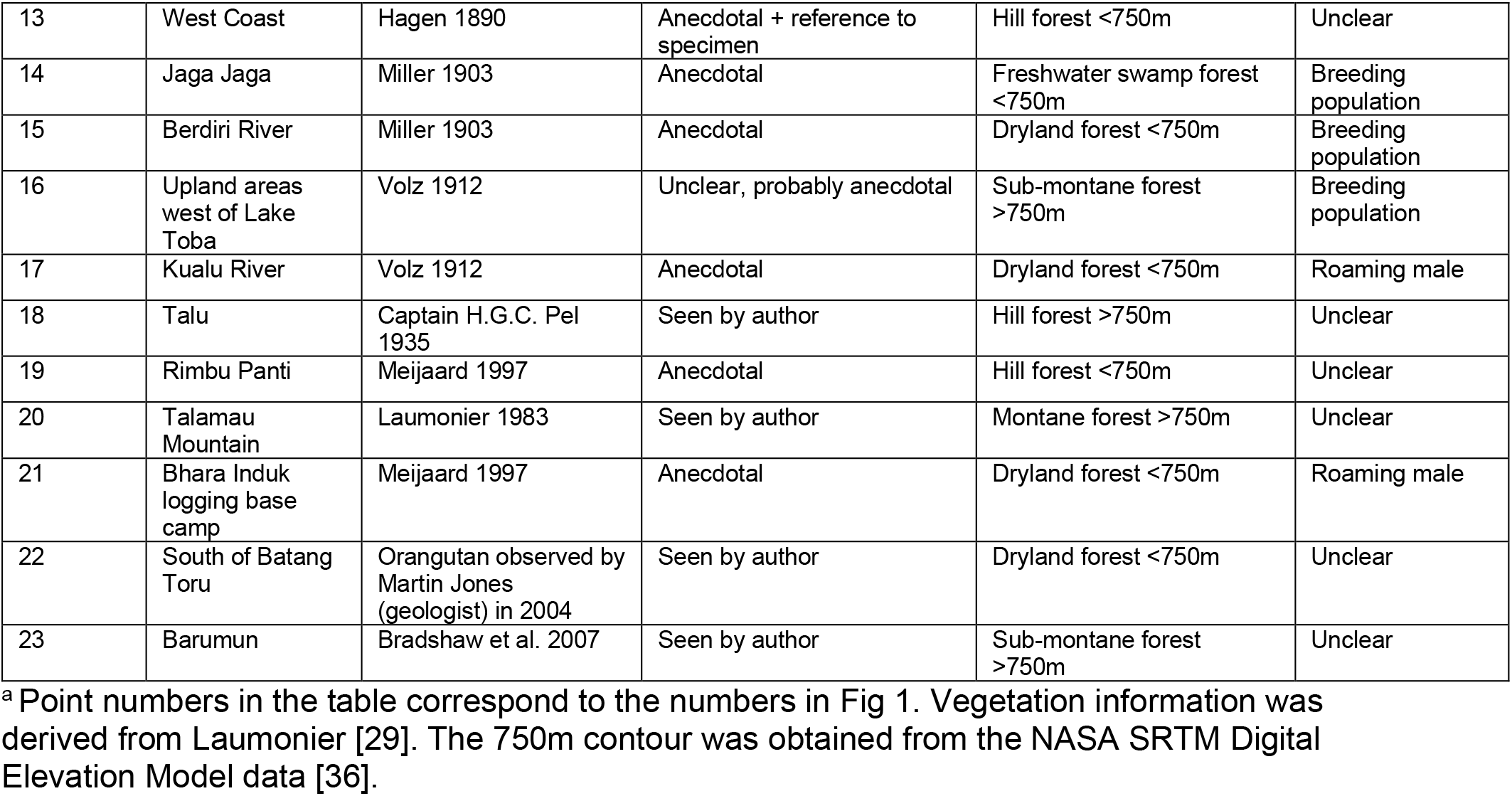
Orangutan records, most likely *P. tapanuliensis*, but outside the current range, that we consider to be reliable.

| Number | Location | Source | Information type | Vegetation/Elevation | Likely status |
| --- | --- | --- | --- | --- | --- |
| 1<sup>a</sup> | Angola | Tulp 1641 | Specimen seen by author | Hill forest <750m | Breeding population |
| 2<sup>a</sup> | Kampar Regency | Müller 1837 | Anecdotal | Dryland forest <750m | Unclear |
| 3 | Around Bukittinggi | Anonymous 1842 | Anecdotal | Hill forest <750m | Unclear |
| 4 | Between Fort de Kock and Bonjol | Anonymous 1842 | Anecdotal | Hill forest <750m | Unclear |
| 5 | Bukit Gedang | Ludeking 1862 | Anecdotal | Sub-montane forest >750m | Roaming male |
| 6 | Tapanoeli | Von Rosenberg 1878 | Anecdotal or seen by author | Dryland forest <750m | Breeding population |
| 7 | Sayur Matinggi | Kramm 1879 | Seen by author | Hill forest >750m | Breeding population |
| 8 | Loemoet | Kramm 1879 | Seen by author | Tall Peat swamp forest <750m | Breeding population |
| 8 | Lumut | Nests observed by Serge Wich and Tine Geurts in June 2001 | Seen by author | Tall Peat Swamp forest <750m | Breeding population |
| 8 | Nests near Lumut | Nests observed by Serge Wich and Tine Geurts in June 2001 | Seen by author | Tall Peat Swamp forest <750m | Breeding population |
| 9 | Hadjoran | Neumann 1885 | Anecdotal | Hill forest >750m | Roaming male |
| 10 | Batang Nilo | L.H. 1888 | Anecdotal | Freshwater swamp forest <750m | Unclear |
| 11 | Lake Maninjau | Twiss 1890 | Anecdotal | Hill forest >750m | Unclear |
| 12 | Bukit Silajang | Twiss 1890 | Anecdotal | Sub-montane forest >750m | Unclear |
| 13 | West Coast | Hagen 1890 | Anecdotal + reference to specimen | Hill forest <750m | Unclear |
| 14 | Jaga Jaga | Miller 1903 | Anecdotal | Freshwater swamp forest <750m | Breeding population |
| 15 | Berdiri River | Miller 1903 | Anecdotal | Dryland forest <750m | Breeding population |
| 16 | Upland areas west of Lake Toba | Volz 1912 | Unclear, probably anecdotal | Sub-montane forest >750m | Breeding population |
| 17 | Kualu River | Volz 1912 | Anecdotal | Dryland forest <750m | Roaming male |
| 18 | Talu | Captain H.G.C. Pel 1935 | Seen by author | Hill forest >750m | Unclear |
| 19 | Rimbu Panti | Meijaard 1997 | Anecdotal | Hill forest <750m | Unclear |
| 20 | Talamau Mountain | Laumonier 1983 | Seen by author | Montane forest >750m | Unclear |
| 21 | Bhara Induk logging base camp | Meijaard 1997 | Anecdotal | Dryland forest <750m | Roaming male |
| 22 | South of Batang Toru | Orangutan observed by Martin Jones (geologist) in 2004 | Seen by author | Dryland forest <750m | Unclear |
| 23 | Barumun | Bradshaw et al. 2007 | Seen by author | Sub-montane forest >750m | Unclear |

<sup>a</sup> Point numbers in the table correspond to the numbers in Fig 1. Vegetation information was derived from Laumonier [29]. The 750m contour was obtained from the NASA SRTM Digital Elevation Model data [36].

### Mapping the historical range

Akçakaya et al. [72] proposed two potential temporal benchmarks for setting historical baselines for species distribution: the year 1500 AD and the year 1750 AD. Species that became extinct before 1500 AD are not assessed for the IUCN Red List, so choosing 1500 AD is a natural link between IUCN Red List and Green Status frameworks [9]. Sanderson [73], however, proposed more flexible dating depending on when locally modern humans started to negatively impact a particular species. For orangutans on Sumatra that could be at least 30,000 years ago [11], but the fossil record from that era is limited to a few sites only and does not allow reliable mapping of the orangutan’s past range, or whether it indeed was the range of *P. tapanuliensis* or *P. abelii*. There is also no reliable information available in our current data for *P. tapanuliensis* in either 1500 AD or 1750 AD, so we choose to use 1890 AD as a time when there are various records that indicate the presence of orangutans through direct observation by the source [i.e., 47, 53, 56] (Fig 3). We also present an alternative historical baseline based on the distribution map drawn by van Heurn in 1935 [65], with additional records from our own dataset (Fig 3).

**Fig 3.**
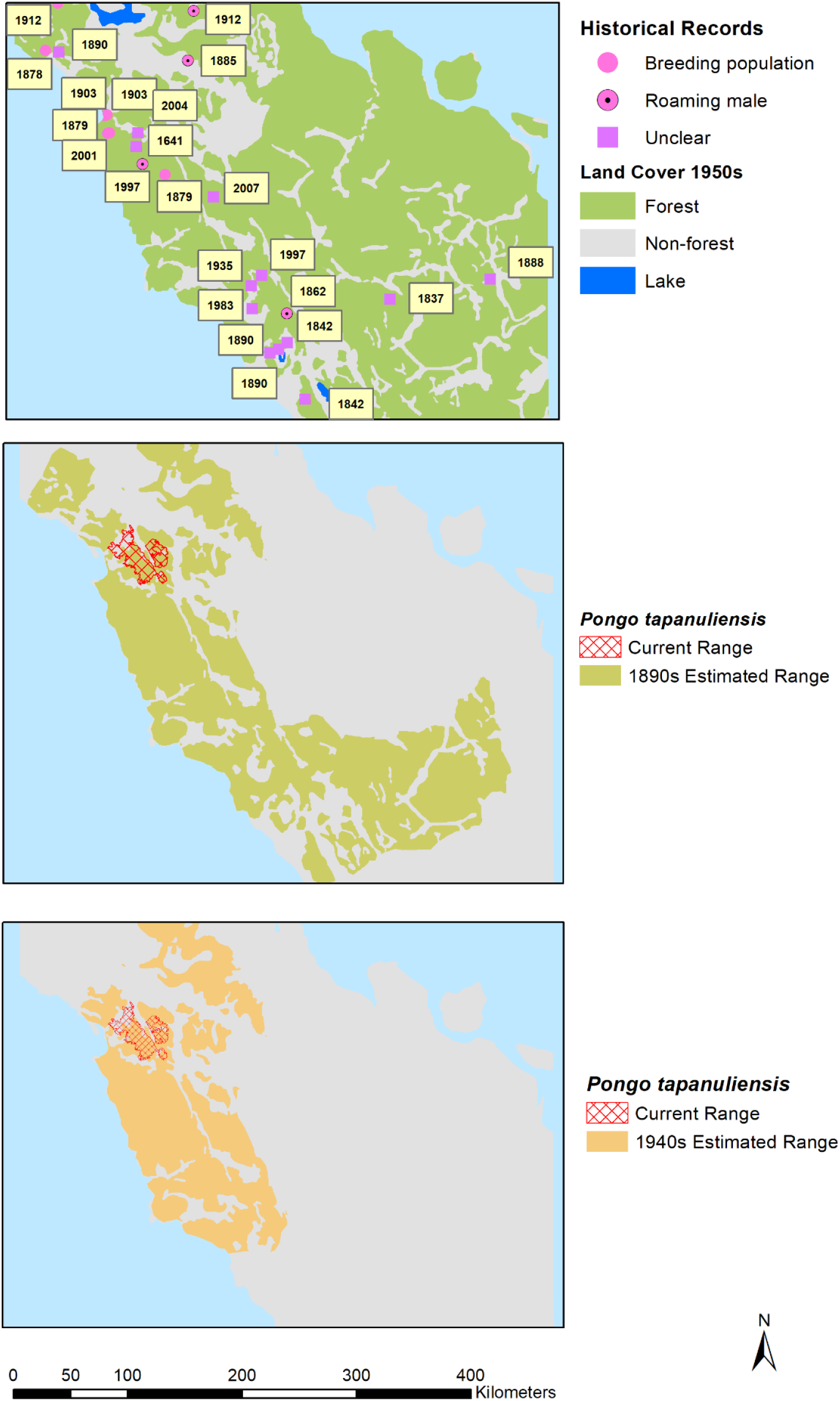
Top panel: Years of historic orangutan records. Middle panel: Estimated historic range in 1890s. Bottom panel: Estimated historic range in 1940s. Reprinted from [26] under a CC BY license, with permission from PT Balai Pustaka, original copyright 1950.

## Discussion

### The historical range of *P. tapanuliensis*

Our historical ecology analysis of *P. tapanuliensis* indicates that the species occurred beyond its current range until quite recently, and has rapidly declined in the past 100 to 150 years. Breeding populations occurred in the Batang Gadis area (Fig 1), probably through much of today’s South Tapanuli and Mandailing Natal Regencies, and further south-east in the Kampar River area. The historical records to the north of the current *P. tapanuliensis* range make it difficult to judge whether these are part of *P. abelii* or *P. tapanuliensis*. For example, the records of orangutans west of Lake Toba, could also refer to populations that still occur in the Pakpak Bharat Regency [74] and the Batu Ardan and Siranggas forest blocks in the Dairi Regency [75], which genetically are closer to *P. abelii* than *P. tapanuliensis* [76]. It is not clear whether the records in the Padang Highlands, Rimbu Panti and Padang Lawas referred to itinerant males or breeding populations but the scarcity of records could indicate that breeding populations became extinct there earlier.

To what extent do these historical data allow us to accurately determine the former range of *P. tapanuliensis*. We know from evidence of Late Pleistocene orangutan fossils in the Padang area (Lida Ajer, Ngalau Sampit, Ngalau Gupin) [77], that orangutans lived in this part of Sumatra at least until some 50,000 years ago. What we do not know is whether this was *P. abelii, P. tapanuliensis*, or a species different from both, as suggested by Drawhorn [78]. No specimens of *P. tapanuliensis* were collected by any of the historical sources, except those reported by Hagen [57] and Beccari [59], but these have not yet been genetically analysed. There is therefore no robust evidence as to whether the orangutans reported from outside the current *P. tapanuliensis* range were *P. tapanuliensis* or *P. abelii*. Further genetic study of the specimens reportedly originating from Padang (RMNH.MAM.544) and Palembang (MZUF-12), and also of fossil teeth from the Padang Caves area (e.g., through proteomic analysis) could shed light on the taxonomic status of the orangutans in Central Sumatra, and their relationship to *P. tapanuliensis*. Based on distribution range patterns, with *P. abelii* clearly restricted to the northern parts of Sumatra [12, 70], however, we consider it most likely that all historical orangutan populations south and south-east of the current range of *P. tapanuliensis* were also *P. tapanuliensis*. If correct, this would indicate a historical extent of occurrence [79] of about 40,796 km^2^ in the 1890s, and about 21,313 km^2^ in the 1940s. If we compare this to the current distribution range of 1,023 km^2^ [14, 15], it suggests that the species currently retains some 2.5% of the 1890s range and 5.0% of the 1940s range. This range included lowland swamp and dry land forests as well as higher elevation forests, suggesting that *P. tapanuliensis* used to occur across a wide range of habitats and is not an ecological specialist of higher elevation areas. It might even be possible to map the range going back further in time, e.g., 1750 AD as recommended by Grace et al. [9]. As populations dwindled and encounters with orangutans became increasingly rare, this may have resulted in the folk zoology regarding mythical creatures (*orang-pendek, gugu, sedapa* etc.) [41, 42] in Bengkulu, Jambi, and South Sumatra, indicating an even larger range. We leave it to other conservation stakeholders to determine which historical range is most appropriate (and reliable) for use as a historical baseline for *P. tapanuliensis*, and also for setting an aspirational recovery target for the long-term future (e.g., 100 years from now) towards full ecological functionality of the species [9, 30, 72].

### Possible drivers of historical declines

While our findings indicate that orangutans disappeared from much of their historical range, it is less clear why the species declined and became locally extinct. Some populations such as those in Lumut, seen by Neumann in 1885 and by SW in 2001, but not seen since, may have become extinct quite recently because of forest loss. Other populations likely disappeared sometime in the 20^th^ century. There have been no recent confirmed records from areas west or southwest of Lake Toba, nor from the Batang Gadis region, south of the current populations. Also, while there are a number of alleged records from the area east of Padang, there are no confirmed recent records. It thus appears that a lot of these populations disappeared around a time when forests were still extensive and the commercial exploitation of forest for timber (starting in the 1970s) or their conversion to plantations (starting in the 1990s) had not yet decimated available habitats. Nevertheless, there had been significant historical deforestation prior to 1950 as shown in Fig 2, mostly for small-holder agriculture and livestock, firewood and timber, and as result of wars and fires [80, 81]. For example, the colonial-era district of *Tapanoeli* (now North, South and Central Tapanuli) had an estimated forest cover of 19,000 km^2^ in a total area of 39,481 km^2^ (i.e., 48% forested) in the 1930s [80]. In 1824, one of the first European visitors to the region was astonished to see that “the plain [north of Batang Toru] was surrounded by hills from five hundred to one thousand feet high, in a state of cultivation; and the whole surrounding country was perfectly free from wood, except the summits of two or three mountains” [82]. Some orangutan populations therefore appear to have become isolated in historical times, when early agricultural development created large grassland areas. So, why did these populations become extinct? This appears to have been a combination of habitat loss and population fragmentation, and mortality rates that exceeded reproductive replacement rates.

Several authors have suggested that orangutan density and range on both Borneo and Sumatra were primarily determined by the ability of people to access areas and hunt orangutans [83, 84]. For example, Jentink (1889) writes that orangutans in Sumatra are only common in swamp areas like those in Singkil which are so inaccessible that they are rarely “stepped on by human feet”, apparently quoting von Rosenberg [51], who had made a similar statement a decade earlier. Wallace (1869) similarly argued that orangutans were common in swamp forest, not because these were particularly suitable ecologically but rather because human hunters rarely went there. Such hunting was certainly common in the orangutan’s range in Sumatra. Schneider [85], for example, writes that Batak people hunt orangutans with blow pipe, spears or shotguns, while young animals are often caught and sold to plantation owners.

Batang Gadis was populated by Loeboe people [86, p. 327] who were nomadic tribes that also occurred in Padang Lawas and “Groot-Mandailing” [87-89]. Another nomadic tribe, the Oeloes occurred around Muara Si Pongi and Pahantan (now Pakantan) [87, 90]. Similar to other nomadic people such as the Kubu further south in Sumatra [91, 92] or the Punan of Borneo [93, 94], nomadic people often prefer primate meat over other meat sources. Kreemer [89] mentioned that the Loeboe people consider primate meat a delicacy. They hunted primates, including Siamang *Symphalangus syndactylus* with blowpipes, and used snares for pigs and deer. Still, there are, to our knowledge, no specific accounts of people from the historical range of *P. tapanuliensis* hunting and eating orangutans. Nevertheless, we consider it likely that, similar to Borneo, orangutans would have been hunted for food. Van den Burg [95], in a general account about orangutans, describes how generally orangutans were shot with poison darts, after which they fell out of the trees and were killed with spears. Alternatively, they were caught alive and killed later. The whitish meat was generally grilled over a fire, and was described as soft and sweet [95]. This is also suggested by the use in local language of *juhut bontar*, or white meat, to describe orangutan [96], while descriptions of sweet meat were similarly recorded by EM on Borneo [25]. Orangutan fat, especially from adult males, was often saved for later use in the preparation of other dishes [95]. Orangutan skins and teeth were further used as amulets on Sumatra, where people hunted them with blowpipes, spears or shotguns, although this account describes the meat as tasting “unpleasant, off-putting and bitter” [85]. We do not know the extent to which hunting and collecting for zoos by foreigners contributed to *P. tapanuliensis* population decline, as was likely the case for *P. abelii* given the large number of animals that were killed and collected [97]. In conclusion, it seems likely that *P. tapanuliensis* were hunted as part of people’s normal selection of big game animals, although whether off-take levels were unsustainable remains unclear.

Marshall et al. [98] examined population viability models with 1%, 2%, and 3% additional mortality in all age classes, running 500 iterations with populations of 250 orangutans. In the best quality orangutan habitats, i.e., mosaic landscapes of swamp, riparian and hill forests [25], annual hunting rates of 1% did not cause population extinction, but did decrease population size. In less-than-optimal habitats, e.g., forests at higher elevation, a 1% level of hunting caused declines to extinction irrespective of initial population size. Higher rates of hunting were unsustainable even in the highest quality habitats [98]. These models were conducted for *P. pygmaeus*, but the authors thought that hunting effects would be similar for Sumatran orangutans. The best orangutan habitats would like be those with the highest soil fertility, which at levels of intermediate rainfall would also be the best areas for agriculture [99]. It is thus likely that historically *P. tapanuliensis* occurred in suboptimal habitats, where the removal of one animal from a population of 100 per year, would drive such a population to extinction. Orangutans are especially vulnerable to the combined effects of fragmentation and unsustainable mortality rates, because females are strongly philopatric and will not move far from their natal ranges [100, 101]. If females are targeted by hunters, this depletes the populations without compensation from female dispersal from nearby forest fragments.

Given the available information, we consider it likely that *P. tapanuliensis* was hunted to extinction in the increasingly fragmented parts of its former range and only survived in the remote and rugged Batang Toru mountains which may have provided orangutans with a refuge from hunting. At the same time, we recognize that in the complex socio-ecological system from which orangutans disappeared many processes may have contributed, and simple, linear cause-effect reasoning may not apply.

### The biases and constraints of a historical perspective

While the use of historical data provides useful insights about the likely historical range of orangutans on Sumatra, and possible drivers of their decline and local extinction, there is uncertainty in the data. Few of the records are based on specimens that provide evidence for the veracity of claims, and some records are only based on hearsay or alleged sightings of orangutans without further evidence. While we critically examined each record, there is subjectivity in interpreting their reliability. For example, we decided not to incorporate the many records of *orang-pendek* in our analysis, even though some of them could well have referred to orangutans. Then again, we did accept sources that reported orangutans seen by local people but not the source, e.g., Neumann [54]. Had the people in Hadjoran mentioned “*orang-pendek*” rather than orangutan, we would have rejected the information, even though it would have related to the same animal. The potential bias of this approach is obvious.

Another form of bias in our study is the literature accessible to us in this study. Most of our information sources are colonial-era explorers, naturalists and hunters, for which we were able to find information in the large numbers of books, newspapers and journals that have been digitized and can be searched and accessed online. This means that we are missing out on two potentially valuable data sources: 1) Local vernacular zoology about orangutans among people that live in the orangutan’s historical range; and 2) Post-independence publications and grey literature from government (e.g., forest inventories), universities (e.g., student survey reports), companies (environmental impact assessments), and local media. There is much vernacular information about orangutan from Borneo (especially Sarawak), but such information does not seem to have been recorded in the anthropological literature for Sumatra. Post-independence writings from the 1950s to ca. 1980s are likely to contain many references to records of orangutans from their historical range, but such information has not yet been captured electronically and remains beyond our reach.

Future studies that would include more local and socio-culturally specific information, would put conservationists in a better position (with the help of local experts, anthropologists, etc.) to understand local drivers of extinction and formulate more targeted interventions. For example, information from indigenous groups that hunt and consume orangutans, versus conflict-related killing of orangutans, versus Muslim taboos against eating orangutans, versus groups that may or may not have specific ritual relations with orangutans [102], could result in locally specific management strategies for reducing killing, harming and capture of orangutans. There are also contextual specificities, e.g., transmigrants from Java or tsunami refugees from Nias Island having very different experiences of forest life, land rights and reactions to orangutans compared to indigenous people in the orangutan’s range. All these nuances relevant to species management require that we go beyond the confines of the data sources used for the current study. There is thus value in the historical ecology approach but there are also limitations, or in the words of the statistician George Box “All models are wrong, but some are useful” [103]. We hope our historic models add useful information to the conservation debate regarding *P. tapanuliensis*.

### Implications for species conservation

What do our findings mean for conservation? The remaining three subpopulations of *P. tapanuliensis* are in apparent decline, threatened by conflict killing and hunting, and loss of lowland habitat [14, 21, 22, 104]. Our insights from past population declines, driven by habitat loss and fragmentation and probably unsustainable mortality rates, indicate that without preventing further losses to the population, even if in the single numbers per year, the last remaining populations of the species are doomed to rapidly decline within several orangutan generation lengths [estimated at 25 years, 23]. Current killing or removal rates of *P. tapanuliensis* already meet or exceed this threshold. Two wild-captured infant *P. tapanuliensis* were reported thus far in 2020, with one confiscated from the owner and the other illegally released to avoid legal repercussions [105, 106]. Three additional infants were confiscated from Tapanuli Selatan, two in 2008 and one in 2015, and one 6-year old female was confiscated from Tapanuli Utara in 2012 (Sumatran Orangutan Conservation Programme pers. comm.). Obtaining wild orangutan infants necessitates killing the mother in nearly all cases [25, 107], hence these infants are assumed to represent two adult females killed in the first six months of 2020 alone, and another three between 2008 and 2015. Such records are indicative of a lowest minimum number of killings, as they represent only criminal acts that have been detected and acted upon, which is a fraction of the total orangutan-related wildlife crime [108, 109]. Records of an adult male killed in 2013 (OIC pers. comm. 2020), a male severely injured by humans in 2019 [110], and another male captured and translocated twice in the past 12 months due to complaints about crop raiding from local community members [111] suggest that killings have been ongoing in recent years, although prior to 2017 most detected incidents would have been recorded as *P. abelii*. While translocation has been used as a response to orangutans in conflict with humans, translocated animals are not monitored beyond a few days following release, sometimes not at all, and their long-term survival is not known. Behavioural traits of female site fidelity and male territoriality, and adaption issues *P. pygmaeus* released in unfamiliar habitats indicate that translocation risks are high and survival rates may be low [108, 112].

Long-term protection of *P. tapanuliensis* requires that mortality rates of <1% per year are maintained over long (decadal) time frames across the species’ range. This also means that that all subpopulations have to remain connected, because once connections between populations are lost this should result in higher extinction risks for the remaining subpopulations, as was modeled for *P. pygmaeus* [35]. Within the subpopulations, the prevention of killing and translocation or rescues is urgently needed, which requires innovative management of crop conflicts [113, 114], and effective law enforcement and awareness campaigns. Such campaigns have so far had insufficient impact on reducing orangutan losses and new approaches may be required [115]. This could include, for example, direct conditional payments to rural communities for maintaining habitats and preventing any deaths or harm, i.e., orangutan guardians [116] or support for “buffer gardens” to concentrate crop losses from orangutan foraging into areas acceptable to communities [117]. Viable conservation solutions that prevent the extinction of *P. tapanuliensis* require an awareness of the specific problem posed by small-scale anthropogenic factors that have driven historical declines. Addressing these factors requires more targeted interventions, for example, through a conservation plan that is tailored specifically to the needs and characteristics of *P. tapanuliensis* and the different socio-ecological drivers of its decline, rather than a generic national-level approach that encompasses a huge range of contexts and all three species [118].

Currently, *P. tapanuliensis* is rated Critically Endangered A4bcd on the IUCN Red List [119] an “observed, estimated, inferred, projected or suspected population size reduction of ≥80% over three generation periods (i.e., 75 years), where the time period must include both the past and the future, and where the reduction or its causes may not have ceased or may not be understood or may not be reversible, based on (b) an index of abundance appropriate to the taxon; (c) a decline in area of occupancy, extent of occurrence and/or quality of habitat; and (d) actual or potential levels of exploitation [23]. The information from the current information makes it likely that a similar decline population size reduction of ≥80% has occurred over the past 75 years, based on the estimated reduction of the Extent of Occurrence [see 79] of 95–97.5 % over 100 to 150 years. This would qualify the species as Critically Endangered A4bcd and A2cd.

Given the high extinction risks, it is important that a comprehensive plan of action is developed for the species that accurately determines how many animals remain, the level of gene flow between subpopulations, current loss rates (including removal of animals in rescues and translocations), and works towards full and permanent protection of all remaining habitat and enforcement of zero unnatural losses. Once remaining populations are secure and viable, the historic range data provide information for considering expansion of the current range to parts of the historic range where the species is now extinct. Such aspirational recovery plans would also ensure that the species once again uses the full range of vegetation types it have evolved in, restoring ecological functionality. Short and long-term conservation plans would need clarity about funding, organizational responsibilities, and a clear, science basis to allow the *P. tapanuliensis* population to stop declining, or better, increase to safer population numbers.

Without such concerted and coordinated action, the remaining populations of *P. tapanuliensis* are doomed to follow their historical predecessors on their path to rapid extinction.

## Acknowledgements

We thank Herman Rijksen for inspiring us to access the historical literature on orangutans, Gabriella Fredriksson, Graham Usher, Gregory Forth, and Liana Chua for providing comments on an earlier version of this manuscript, Matthew Minarchek for sharing Carpenter’s interview data, which allowed us to update the historical range maps, Molly Grace and Mike Hoffmann for suggesting how we could map the historical ranges, and Yves Laumonier for granting permission to use the Vegetation of Sumatra digital maps, and Wiwit Siswarini and Bart van Assen for helping to obtain copyright permission from PT Balai Pustaka. We also thank three reviewers for their constructive comments.

